# Recommendations for improving statistical inference in population genomics

**DOI:** 10.1101/2021.10.27.466171

**Authors:** Parul Johri, Charles F. Aquadro, Mark Beaumont, Brian Charlesworth, Laurent Excoffier, Adam Eyre-Walker, Peter D. Keightley, Michael Lynch, Gil McVean, Bret A. Payseur, Susanne P. Pfeifer, Wolfgang Stephan, Jeffrey D. Jensen

**Affiliations:** School of Life Sciences, Arizona State University, Tempe, US; Department of Molecular Biology and Genetics, Cornell University, Ithaca, US; School of Biological Sciences, University of Bristol, Bristol, UK; Institute of Evolutionary Biology, School of Biological Sciences, University of Edinburgh, Edinburgh, UK; Institute of Ecology and Evolution, University of Berne, Berne, CH; School of Life Sciences, University of Sussex, Brighton, UK; Institute of Ecology and Evolution, School of Biological Sciences, University of Edinburgh, Edinburgh, UK; Big Data Institute, Li Ka Shing Centre for Health Information and Discovery, University of Oxford, Oxford, UK; Laboratory of Genetics, University of Wisconsin-Madison, Madison, US; Natural History Museum, Berlin, DE

**Keywords:** population genetics, population genomics, statistical inference, model-fitting, positive selection, purifying selection, background selection, selective sweeps, genome scans, demography

## Abstract

The field of population genomics has grown rapidly in response to the recent advent of affordable, large-scale sequencing technologies. As opposed to the situation during the majority of the 20th century, in which the development of theoretical and statistical population-genetic insights out-paced the generation of data to which they could be applied, genomic data are now being produced at a far greater rate than they can be meaningfully analyzed and interpreted. With this wealth of data has come a tendency to focus on fitting specific (and often rather idiosyncratic) models to data, at the expense of a careful exploration of the range of possible underlying evolutionary processes. For example, the approach of directly investigating models of adaptive evolution in each newly sequenced population or species often neglects the fact that a thorough characterization of ubiquitous non-adaptive processes is a prerequisite for accurate inference. We here describe the perils of these tendencies, present our consensus views on current best practices in population genomic data analysis, and highlight areas of statistical inference and theory that are in need of further attention. Thereby, we argue for the importance of defining a biologically relevant baseline model tuned to the details of each new analysis, of skepticism and scrutiny in interpreting model-fitting results, and of carefully defining addressable hypotheses and underlying uncertainties.

## Introduction

### A brief overview

Population genomic inference – the use of data on molecular variation within species and divergence between species to infer evolutionary processes – has become widely embraced and highly utilized in fields including evolutionary biology, ecology, anthropology, agriculture, and medicine. The underlying questions may be demographic in nature, be it estimating the timing of the peopling of the world [1] or of viral transmission in a congenitally infected newborn [2]; alternatively, they may concern the selective history of specific populations, be it identifying mutations that confer cryptic coloration in species adapting to major post-glacial climatic and geological changes [3] or viral drug-resistance to clinical therapeutics [4].

The foundational work allowing for the dissection of these evolutionary processes from levels and patterns of variation and divergence was conducted by Fisher, Wright, and Haldane nearly a century ago (*e.g*., [5–7]; for a historical overview, see [8]). This work demonstrated the possibility of studying evolution at the genetic level, integrating the revolutionary ideas of Darwin [9] with the turn-of-the-century appreciation of Mendel’s [10] research. However, as was famously described by Lewontin [11], this initial theoretical progress during the first half of the 20th century was “like a complex and exquisite machine, designed to process a raw material that no one had succeeded in mining”. With the first ‘mining’ of population-level molecular variation in the 1960s (see [12]), this machine was put to work. The next major steps forward were provided by Kimura and Ohta, who offered a comprehensive framework for studying DNA and protein sequence variation based on these fundamental theoretical insights – the Neutral Theory of Molecular Evolution [13–15] – an advance for which molecular biology also provided support [16]. Despite some claims to the contrary [17], Kimura and Ohta’s initial postulates have since been largely validated [18–19], and have provided a means to interpret observed molecular variation and divergence within the context of constantly occurring evolutionary processes including mutation, genetic drift, and purifying selection. While ascribing an important role for positive selection at the level of phenotypic evolution (consistent with Darwin’s initial notions), the Neutral Theory hypothesizes that at the genetic level beneficial mutations are rare compared to the much larger input of neutral, nearly neutral, and deleterious mutations that are constantly raining down on the genomes of all species. Accordingly, episodes of positive selection per nucleotide are rare compared to genetic drift and purifying selection. However, the significant effects on evolution at linked sites caused by fitness-altering mutations have been described in detail in the decades since Kimura’s initial formulation of the Neutral Theory [20–22].

With this framework and the availability of datasets to which it could be applied, statistical approaches for analyzing molecular data began to proliferate, frequently employing some form of neutral expectation as a null model. A wide range of rather sophisticated statistical machinery is now available for reconstructing histories of population size change, population subdivision and migration (*e.g.,* [23–24]), for identifying beneficial mutations based on patterns associated with selective sweeps (*e.g.,* [25–26]), for quantifying the distribution of fitness effects (DFE) of newly arising mutations (*e.g.,* [27–28]), as well as for estimating rates of mutation (*e.g.,* [29–31]) and recombination (*e.g.,* [32–34]). These approaches operate in a variety of statistical frameworks (see [35–37]), and utilize various aspects of the data – including the frequencies of variants in a sample (the site frequency spectrum, SFS), associations between variants (linkage disequilibrium, LD), and/or levels and patterns of between-species divergence at contrasted site classes (*e.g.,* synonymous versus non-synonymous sites).

### Challenges of model choice and parameter fitting

The growing variety of statistical approaches and associated software implementations presents a dizzying array of choices for any given analysis; although many approaches share the same aims, there also exist important differences. For example, some approaches require a relatively high level of coding ability to implement while others may be applied in easy-to-use software packages; while some are well-tested and justified by population-genetic theory, others are not. Moreover, even the process of translating raw sequencing data into the allele calls and genotypes used as input for these approaches is accompanied by uncertainty that depends on sequencing quality and coverage, availability of a reference genome, and choice of variant calling and filtering strategies [38–39]. Adding to this complexity, it has become increasingly clear that demographic estimation may be highly biased when selection and recombination-associated biased gene conversion are neglected [40–41], whereas estimates of selection intensity and recombination rate may be highly biased when neglecting demographic effects [42–45]. This creates a circular problem when commencing any new analysis: one needs information about the demographic history to estimate parameters of recombination and selection, while at the same time one needs information about recombination and selection to estimate the demographic history. An additional challenge, and a frustration for many, is that there is no single ‘best approach’; the correct analysis tools to use, and indeed which questions can be answered at all, depend entirely on the details of the organism under study [46]. Specifically, biological parameters that vary among species – including evolutionary parameters (*e.g*., effective population size (*N_e_*), mutation rates, recombination rates, and population structure and history), genome structure (*e.g*., the distribution of functional sites along the genome), and life history traits (*e.g*., mating system) – must all be considered in order to define addressable hypotheses and optimal approaches.

Beyond these initial considerations, a more difficult issue often emerges. Namely, very different models may be found to provide a good fit to the observed data (*e.g*., [47]; see [48] for a phylogenetic perspective on the topic). In other words, particular parameter combinations may be found under competing models that are all capable of predicting the observed patterns of variation. For example, assuming neutrality, one may match an empirical observation at a locus by fitting the timing, severity, and duration of a population bottleneck; or, alternatively, when assuming a constant population size, by fitting the rate and mean strength of selective sweeps. This fact alone implies a simple truism: the ability to fit the parameters of one’s preferred model to data does not alone represent proof of biological reality. Rather, it suggests that this model is one – out of potentially very many – that represents a viable hypothesis, which should be further examined via subsequent analyses or experimentation.

Examples abound of enthusiastic promotion of a single preferred model, only to be tempered by subsequent demonstrations of the fit of alternative and often simpler / more biologically realistic models. For example, the view that segregating alleles may be commonly maintained by balancing selection [49] was tempered by the realization that genetic drift is often a sufficient explanation [14], and the view that genome-wide selective sweeps on standing variation are pervasive [50–51] was tempered by the realization that neutral population histories can result in similar patterns [47, 52]. While one may readily find such examples of using episodic or hypothesized processes to fit large-scale data patterns by neglecting to define expectations arising from common and certain-to-be-occurring processes, determining which models to evaluate, and how to interpret the fit of a model and its alternatives, are challenges for all researchers. To better illustrate this point, Figure 1 presents three scenarios (constant population size with background selection, constant population size with background selection and selective sweeps, and a population bottleneck with background selection and selective sweeps), and provides the fit of each of those scenarios to two incorrect models (population size change assuming strict neutrality, and recurrent selective sweeps assuming constant population size). As is shown in the figure, each scenario can be well fit by both incorrect models, with selective sweeps and population bottlenecks generally being confounded, as well as background selection and population growth, as has been described several times before (*e.g*., [40, 53–55]).

**Figure 1.**
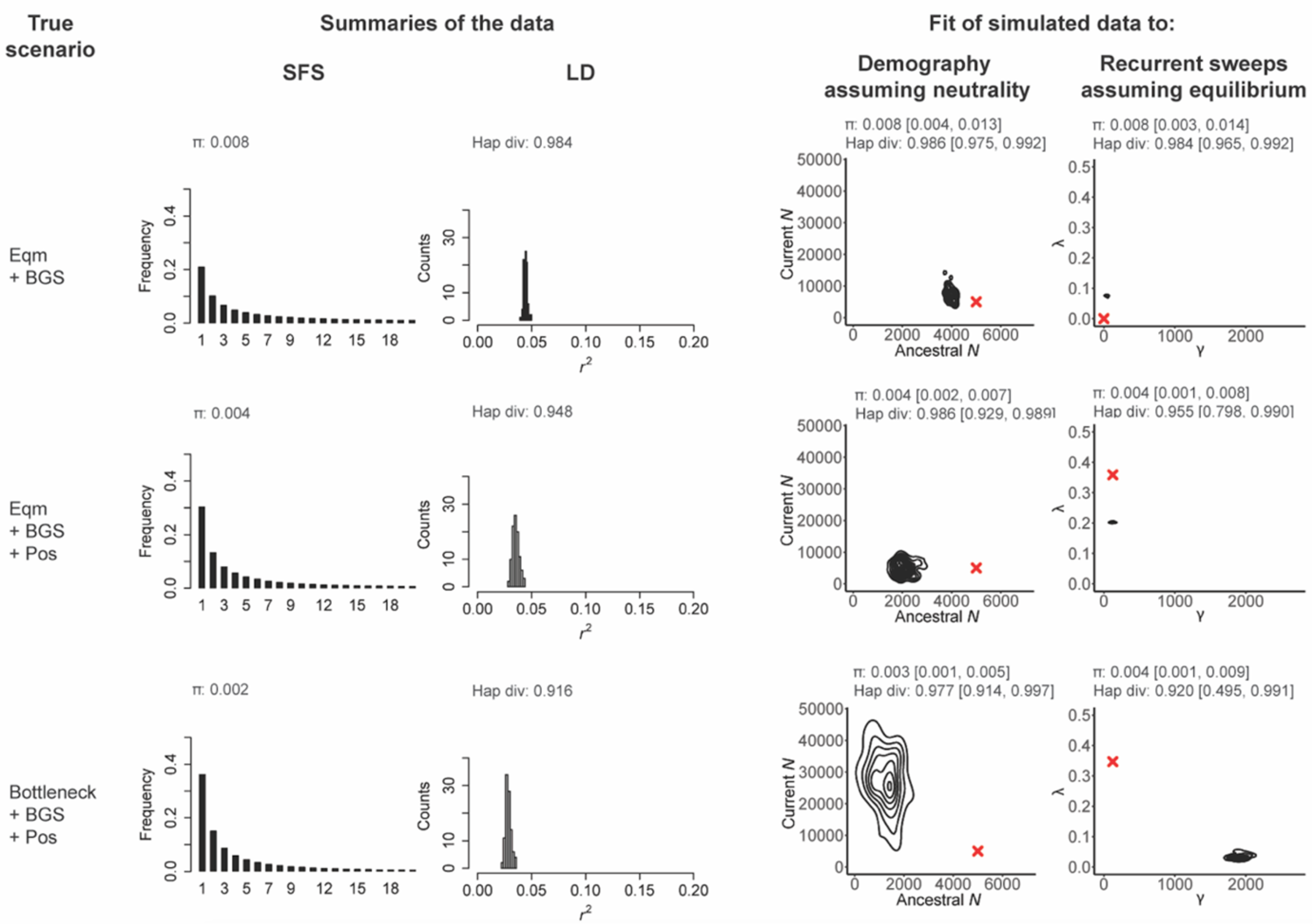
Incorrect models may often readily be fit to a given dataset. Here we present three scenarios varying from simple to more complex: the first row presents a constant-sized population experiencing background selection (denoted by ‘Eqm + BGS’), the second row is the same scenario with the addition of recurrent selective sweeps (denoted by ‘Eqm +BGS + Pos’), and the final row adds a population bottleneck (denoted by ‘Bottleneck + BGS + Pos’). For each scenario, the resulting site frequency spectra (SFS, truncated to *n* = 20) and linkage disequilibrium (*r^2^*) distributions are given, together with mean pairwise (π) and haplotype diversity. To these simulated data we fit two incorrect models; one assuming all sites are neutral but including a change in population size, and a second model in which there are recurrent selective sweeps, no change in population size, and all mutations are assumed to be neutral or beneficial (with a population-scaled beneficial selection coefficient (γ) and the fraction of beneficial substitutions (λ) being estimated from the data). For each inference panel, the red cross gives the true value, the distribution presents the joint-posterior obtained from the ABC analysis, and the summary statistics given above the posteriors represent the mean values, and the range from the 95% CIs, obtained from posterior checks. In all cases, exonic sites (*i.e*., directly selected sites) were masked, and the summary statistic calculations as well as inference is based only on neutral regions (see Methodology). As shown, demographic and selection models can be fit to all datasets, often resulting in strong mis-inference when the assumptions underlying the estimation procedure are violated.

### Methodology

In order to provide a series of examples to accompany key points - as with the above Figure 1 - both forward-in-time simulations and coalescent simulations were performed for (1) the inference of demographic history assuming complete neutrality, (2) the inference of positive selection assuming constant population size, and (3) obtaining test datasets representing different evolutionary scenarios. While any statistical framework involving model / parameter exploration and comparison may be consistent with our recommendations, we here utilize approximate Bayesian computation (ABC) for our examples, as it is a particularly useful framework for quantifying uncertainty and for exploring complex models.

In all simulations, a chromosomal segment of 99,012 bp was simulated with an intron-exon-intergenic structure resembling the *D. melanogaster* genome. Each gene comprised five exons (of 300 bp each) and four introns (of 100 bp each) separated by intergenic regions of length 1,068 bp. Such a construct resulted in a total of 33 genes across the simulated segment.

Population parameters were chosen to resemble those from *D. melanogaster* populations following Campos *et al.* [56], assuming an effective population size (*N*_e_) of 10^6^ individuals with a mean mutation rate (μ) of 4.5 × 10^-9^ per base pair /generation and a mean recombination rate (*r*) of 1 × 10^-8^ per base pair /generation. For computational efficiency, all parameters were rescaled by a factor of 200.

### Modeling and inference of demographic history

A simple demographic history was modeled in which a single population undergoes an instantaneous change from an ancestral size (*N_anc_*) to a current size (*N_cur_*), τ generations ago. Priors for both *N_anc_* and *N_cur_* were sampled from a loguniform distribution between 10 and 50,000, while priors for the time of change (τ) were sampled from a loguniform distribution between 10 and *N_cur_*. One hundred replicates were simulated for each parameter combination. Simulations required for ABC were performed in *msprime* v. 0.7.3 [57] assuming complete neutrality. Mutation and recombination rates were assumed to be constant across the genome and across replicates.

### Modeling and inference of positive selection

A recurrent selective sweep scenario was modeled in which only neutral and beneficial autosomal mutations were allowed, with simulations performed using *SLiM* v. 3.1 [58]. Introns and intergenic regions were assumed to be neutral, while exons experienced beneficial mutations with fitness effects sampled from an exponential distribution with mean *S* for homozygotes, assuming semi-dominance. The two parameters varied were the mean population-scaled strength of selection, γ = 2*N_anc_S*, and the proportion of new beneficial mutations, *f_pos_*. Priors for these parameters were sampled from a loguniform distribution such that γ ∈ [0.1, 10000] and *f_pos_* ∈ [0.00001, 0.01]. For all parameter combinations, the true rate of beneficial substitutions per site (*d_a_*) and the true fraction of substitutions due to beneficial mutations (λ, which is related to the α parameter of Eyre-Walker & Keightley [59]) were calculated using the total number of fixations (as provided by *SLiM*), with λ observed to range from 0-0.85 depending on the underlying parameters. Parameter inference was performed for γ and *d_a_* and the corresponding λ was inferred using 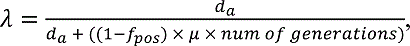 where it was assumed that 1 − *f_pos_*∼1. Populations had a constant size of 5000 diploid individuals with constant mutation and recombination rates, as specified above. Simulations were run for 100,100 generations (*i.e.,* 20*N*_e_ + 100 generations).

### ABC

The sample size was set to 100 haploid genomes (or 50 diploid individuals). Under the demographic and selection models described above, all exonic regions were masked and the mean and variance (across replicates) of the following summary statistics were calculated: number of segregating sites, nucleotide site diversity (π), Watterson’s theta (θ*_W_*), θ*_H_, H′*, Tajima’s *D*, number of singletons, haplotype number and frequency distribution, and statistics summarizing LD (*r^2^, D, D′*). All statistics were calculated in non-overlapping sliding windows of 2 kb using *pylibseq* v. 0.2.3 [60]. ABC was performed with the R package “abc” v. 2.1 [61] using all summary statistics, with “neural net” to account for non-linearity between statistics and parameters. A 100-fold cross-validation was used to identify the optimum tolerance level, which was found to be 0.05 (*i.e.,* 5% of the simulations were accepted during ABC inference to estimate the posterior probability of each parameter). Point estimates of each inferred parameter were calculated as the weighted medians of the posterior estimates.

### Simulations of different evolutionary scenarios as ‘true scenarios’

To consider more biologically realistic models and evaluate model violations, a number of evolutionary scenarios were simulated (using *SLiM*) as follows:

a. Background selection: Exons experienced deleterious mutations modeled by a discrete DFE comprised of four non-overlapping uniform distributions, representing the effectively neutral (−1 < 2*N_anc_*_’_*S* ≤ 0), weakly deleterious (−10 < 2*N_anc_S* ≤ −1), moderately deleterious (−100 < 2*N_anc_S* ≤ −10), and strongly deleterious (2*N_anc_S* ≤ −100) classes of mutations. All four bins were assumed to contribute equally to new mutations (*i.e*., 25% of all new mutations belonged to each class of mutation).
b. Positive selection: Exons experienced beneficial mutations with γ = 125 and *f_pos_* = 2.2 × 10^-3^ (modified from [56]), resulting in λ ≈ 0.35.
c. Population size change: A population decline was simulated such that the population declined from 5000 to 100 individuals instantaneously 100 generations ago. A population expansion was similarly simulated with parameters *N_anc_* = 5000 and *N_cur_* = 10000. A population bottleneck model was also simulated such that *N_anc_* = *N_cur_* = 5000, and a bottleneck occurred 2000 generations ago with a reduction to 1% of the population size for 100 generations.
d. SNP ascertainment: Genotype error was modeled as an inability to detect the true number of singletons when using low-coverage population-genomic data to call variants [38]. To model this scenario, a random set of singletons, representing a third of all singletons present in the sample, were removed.
e. Progeny skew: A skew in the offspring distribution (ψ) was modeled such that 5% and 10% of the population was replaced by the offspring of a single individual each generation ([62], and see [63–64]).
f. Variation in mutation and recombination rates across the genome (*e.g*., [65–67]): Every 10 kb of the ≈100 kb genomic region considered was assumed to have a different mutation and recombination rate. For every simulated replicate, these rates were sampled from a Gaussian distribution with the same mean as above, and a coefficient of variation of 0.5. Negative values were truncated to 0.

### Posterior checks

For the purposes of illustration, an example of posterior checks are provided in Figure 1 (*i.e.,* showing a simple evaluation of the fit of the inferred posteriors under the incorrect models to the true scenarios under consideration). Specifically, the mean estimates of the inferred parameters were used to simulate the “best-fitting model” in *SLiM* v. 3.1 [58]. Exons were masked and summary statistics were calculated as above in windows of 2 kb using *pylibseq* v.0.2.3 [60]. In order to simulate the inferred models of positive selection, *f_pos_* was calculated from λ assuming a Wright-Fisher diploid population of size *N* and a total mutation rate of μ*_tot_* (which for our purpose is the same as μ). Thus, μ*_b_* = *f_pos_* × μ*_tot_* and μ*_neu_* = (1 − *f_pos_*) × μ*_tot_* where μ*_b_* and μ*_neu_* are the beneficial and neutral mutation rates, respectively. Given a value of λ, and assuming that the distribution of fitness effects of beneficial mutations is exponential (with mean ^—^*S*), we calculate *f_pos_* as follows:
given that

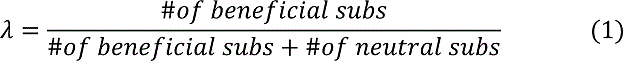

where

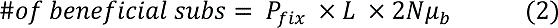

and

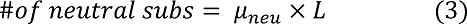

where *L* is the length of the region being considered and *P_fix_* is the probability of fixation of beneficial mutations, given by:

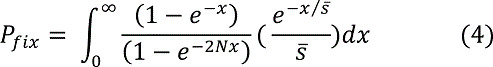

Substituting (2) and (3) in (1), and rearranging we get

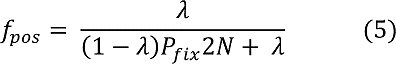

Integrating (4) in *R* and substituting it into (5) gives us values of *f_pos_*.

Statistics were calculated in non-overlapping windows of 2 kb and intervals (CIs) were calculated as the 0.025 and 0.975 quantiles of the distribution of the statistics.

### Recommendations

#### Constructing an appropriate baseline model for population genomic analysis

The somewhat disheartening exercise of fitting incorrect models to data (as depicted in Figure 1) naturally raises the questions of whether, and if so how, accurate evolutionary inferences can be extracted from DNA sequences sampled from a population. The first point of importance is that the starting point for any genomic analysis should be the construction of a biologically relevant baseline model, which includes the processes that must be occurring and shaping levels and patterns of variation and divergence across the genome. This model should include mutation, recombination, and gene conversion (each as applicable), purifying selection acting on functional regions and its effects on linked variants (*i.e.,* background selection [21, 68–69]), as well as genetic drift as modulated by, amongst other things, the demographic history and geographic structure of the population. Depending on the organism of interest, there may be other significant biological components to include, such as mating system, progeny distributions, ploidy, and so on (though, for certain questions of interest, some of these biological factors may simply be included in the resulting effective population size). It is thus helpful to view this baseline model as being built from the ground up for any new data analysis. Importantly, the point is not that these many parameters need to be fully understood in a given population in order to perform any evolutionary inference, but rather that they all require consideration, and that the effects of uncertainties in their underlying values on downstream inference can be quantified.

However, even prior to considering any biological processes, it is important to investigate the data themselves. Firstly, there exists an evolutionary variance associated with the myriad of potential realizations of a stochastic process, as well as the statistical variance introduced by finite sampling. Secondly, it is not advisable to compare one’s empirical observations, which may include missing data, variant calling or genotyping uncertainty (*e.g*., effects of low coverage), masked regions (*e.g*., regions in which variants were omitted due to low mappability and/or callability) and so on, against either an analytical or simulated expectation that lacks those considerations and thus assumes optimal data resolution [70]. The dataset may also involve a certain ascertainment scheme, either for the variants surveyed [71], or given some pre-defined criteria for investigating specific genomic regions (*e.g*., regions representing genomic outliers with respect to a chosen summary statistic [72]). For the sake of illustration, Figure 2 follows the same format as Figure 1, but considers two scenarios: population growth with background selection and selective sweeps, and the same scenario together with data ascertainment (in this case, an under-calling of the singleton class). As can be seen, due to the changing shape of the frequency spectra, neglecting to account for this ascertainment can greatly affect inference, considerably modifying the fit of both the incorrect demographic and incorrect recurrent selective sweep models to the data.

**Figure 2.**
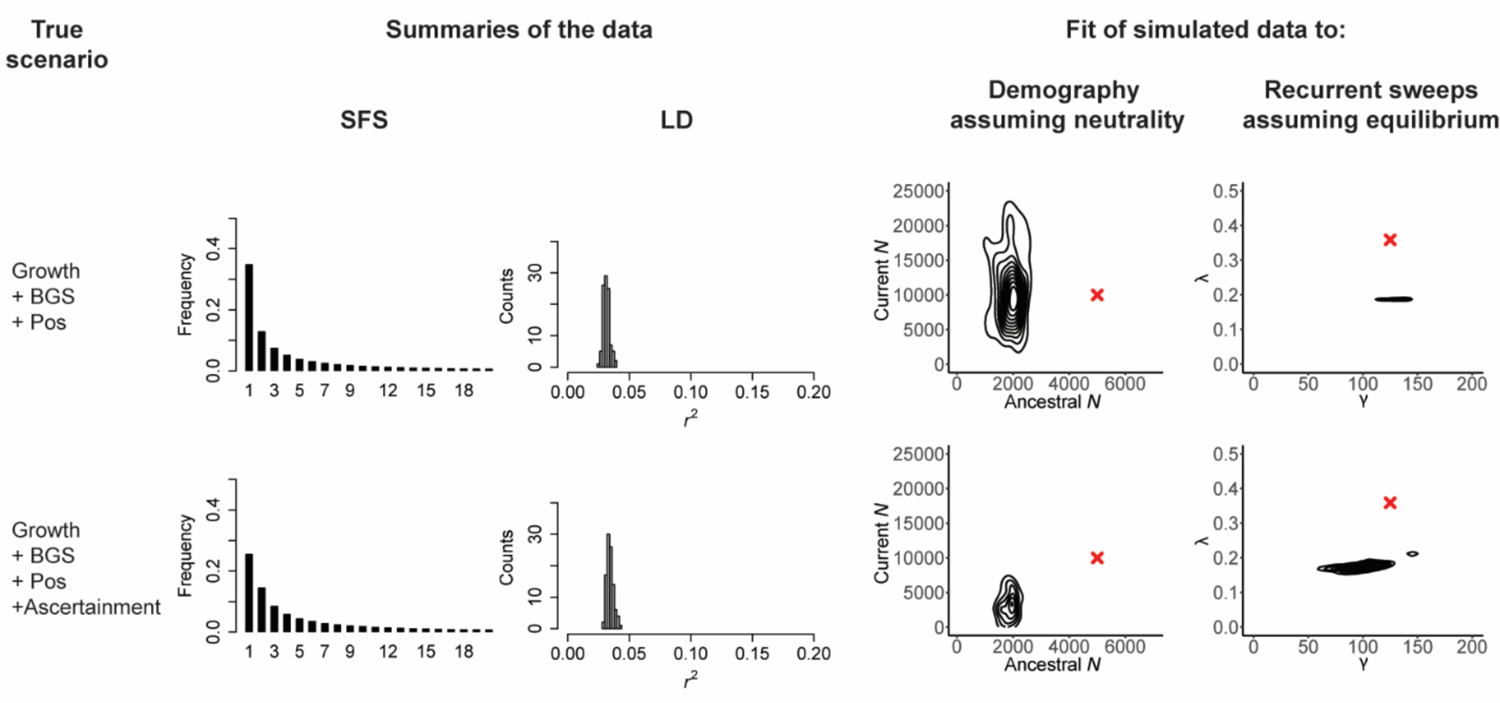
Ascertainment errors may amplify mis-inference, if not corrected. As in Figure 1, the scenarios are given in the first column, here population growth with background selection and recurrent selective sweeps (‘Growth + BGS + Pos’), as well as the same scenario in which the imperfections of the variant-calling processes are taken into account – in this case, one-third of singletons are not called (‘Growth + BGS + Pos + Ascertainment’). The middle columns present the resulting SFS and LD distributions, and the final columns provide the joint posterior distributions when the data are fit to two incorrect models: a demographic model that assumes strict neutrality, and a recurrent selective sweep model that assumes a constant population size. All exonic (*i.e*., directly selected) sites were masked prior to analysis. Red crosses indicate the true values. As shown, unaccounted for ascertainment errors may contribute to mis-inference.

Hence, if sequencing coverage is such that rare mutations are being excluded from analysis, due to an inability to accurately differentiate genuine variants from sequencing errors, the model used for subsequent testing should also ignore these variants. Similarly, if multiple regions are masked in the empirical analysis due to problems such as alignment difficulties, the expected patterns of LD that are observable under any given model may be affected. Furthermore, while the added temporal dimension of time-series data has recently been shown to be helpful for various aspects of population genetic inference [73–76], such data in no way sidestep the need for an appropriate baseline model, but simply requires the development of a baseline that matches the temporal sampling. In sum, as these factors can greatly affect the power of planned analyses and may introduce biases, the precise details of the dataset (*e.g*., region length, extent and location of masked regions, the number of callable sites, and ascertainment) and study design (*e.g.,* sample size and single time-point versus time-series data) should be directly matched in the baseline model construction.

Once these concerns have been satisfied, the first biological addition will logically be the mutation rate and mutational spectrum. For a handful of commonly studied species, both the mean of, and genomic heterogeneity in, mutation rates have been quantified via mutation-accumulation lines and/or pedigree studies [77]. However, even for these species, ascertainment issues remain complicating [78], variation amongst individuals may be substantial [79], and estimates only represent a temporal snapshot of rates and patterns that are probably changing over evolutionary time-scales and may be affected by the environment [31, 80]. In organisms lacking experimental information, often the best available estimates come either from a distantly related species or from molecular clock-based approaches. Apart from stressing the importance of implementing either of the experimental approaches in order to further refine mutation-rate estimates for such a species of interest, it is noteworthy that this uncertainty can also be modeled. Namely, if proper estimation has been performed in a closely related species, one may quantify the expected effect on observed levels of variation and divergence of higher and lower rates. The variation in possible data observations induced by this uncertainty is thus now part of the underlying model.

The same logic follows for the next parameter addition(s): crossing over / gene conversion, as applicable for the species in question. For example, for a subset of species, per-generation crossover rates in cM per Mb have been estimated by comparing genetic maps based on crosses or pedigrees with physical maps [81–83]. In addition, recombination rates scaled by the effective population size have also been estimated from patterns of LD (*e.g*., [84–85]) – though this approach typically requires assumptions about evolutionary processes that may be violated (*e.g.,* [42]). As with mutation, the effects on downstream inference arising from the variety of possible recombination rates – whether estimated for the species of interest or a closely related species – can be modeled.

The next additions to the baseline model construction are generally associated with the greatest uncertainty – the demographic history of the population, and the effects of direct and linked purifying selection. This is a difficult task given the virtually infinite number of potential demographic hypotheses (*e.g*., [86]); furthermore, the interaction of selection with demography is inherently non-trivial and difficult to treat (*e.g*., [55, 87–88]). This realization continues to motivate attempts to jointly estimate the parameters of population history together with the DFE of neutral, nearly neutral, weakly deleterious and strongly deleterious mutations – a distribution which is often estimated in both continuous and discrete forms [89]. One of the first important advances in this area used putatively neutral synonymous sites to estimate changes in population size based on patterns in the SFS and conditioned on that demography to fit a DFE to non-synonymous sites, which presumably experience considerable purifying selection [90–92]. This stepwise approach may become problematic, however, for organisms in which synonymous sites are not themselves neutral [93–95], or when the SFS of synonymous sites is affected by background selection, which is probably the case generally given their close linkage to directly selected non-synonymous sites ([41], and see [96–97]).

In an attempt to address some of these concerns, Johri *et al.* [44] recently developed an ABC approach that relaxes the assumption of synonymous site neutrality and corrects for background selection effects by simultaneously estimating parameters of the DFE alongside population history. The posterior distributions of the parameters estimated by this approach in any given data application (*i.e*., characterizing the uncertainty of inference) represent a logical treatment of population size change and purifying / background selection for the purposes of inclusion within this evolutionarily relevant baseline model. That said, the demographic model in this implementation is highly simplified, and extensions are needed to account for more complex population histories. In particular, estimation biases that may be expected owing to the neglect of cryptic population structure and migration, and indeed the feasibility of co-estimating population size change and the DFE together with population structure and migration within this framework, all remain in need of further investigation. While such simulation-based inference (see [98]), including ABC, provides one promising platform for joint estimation of demographic history and selection, progress on this front has been made using alternative frameworks as well [99–100], and developing analytical expectations under these complex models should remain as the ultimate, if distant, goal. Alternatively, in functionally sparse genomes with sufficiently high rates of recombination, such that assumptions of strict neutrality are viable for some genomic regions, multiple well-performing approaches have been developed for estimating the parameters of much more complex demographic models (*e.g*., [101–104]). In organisms for which such approaches are applicable (*e.g*., certain large, coding sequence sparse vertebrate and land-plant genomes), this intergenic demographic estimation assuming strict neutrality may helpfully be compared to estimates derived from data in or near coding regions that account for the effects of direct and linked purifying selection [41, 44, 105]. For newly studied species lacking functional annotation and information about coding density, following the joint estimation procedure would remain as the more satisfactory strategy in order to account for possible background selection effects.

#### Quantifying uncertainty in model choice and parameter estimation, investigating potential model violations, and defining answerable questions

One of the useful aspects of these types of analyses is the ability to incorporate uncertainty in underlying parameters under relatively complex models, in order to determine the impact of such uncertainty on downstream inference. The computational burden of incorporating variability in mutation and recombination rate estimates, or drawing from the confidence- or credibility-intervals of demographic or DFE parameters, can be met with multiple highly flexible simulation tools [58, 106–107]. These are also useful programs for investigating potential model violations that may be of consequence. For example, if a given analysis for detecting population structure assumes an absence of gene flow, it is possible to begin with one’s constructed baseline model, add migration parameters to the model in order to determine the effects of varying rates and directions of migration on the summary statistics being utilized in the empirical analysis, and thereby quantify how a violation of that assumption may affect the subsequent conclusions.

Similarly, if an analysis assumes the Kingman coalescent (*e.g*., a small progeny distribution such that at most one coalescent event occurs per generation), but the organism in question could violate this assumption (*i.e*., with the large progeny number distributions associated with many plants, viruses, and marine spawners, or simply owing to the relatively wide array of evolutionary processes that may similarly lead to multiple merger coalescent events), these distributions may too be modeled in order to quantify potential downstream mis-inference.

To illustrate this point, Figure 3 considers two scenarios of constant population size and strict neutrality but with differing degrees of progeny skew, to demonstrate that a violation of this sort that is not corrected for may result in severely under-estimated population sizes as well as the false inference of high rates of strong selective sweeps. In this case, the mis-inference arises from the reduction in contributing ancestors under these models, as well as to the fact that neutral progeny skew and selective sweeps can both generate multiple-merger events [63-64, 108-109]. Similarly, one may investigate the assumptions of constant mutation or recombination rates when they are in reality variable. As shown in Figure 4, when these rates are assumed to be constant as is common practice, but in reality vary across the genomic region under investigation, the fit of the (incorrect) demographic and selection models considered may again be substantially modified. Notably, this rate heterogeneity may inflate the inferred strength of selective sweeps. While Figures 3 and 4 serve as examples, the same investigations may be made for cases such as a fixed selective effect when there is in reality a distribution, independent neutral variants when there is in reality linkage disequilibrium, panmixia when there is in reality population structure, and so on. Simply put, even if a particular biological process / parameter is not being directly estimated, its consequences can nonetheless be explored.

**Figure 3.**
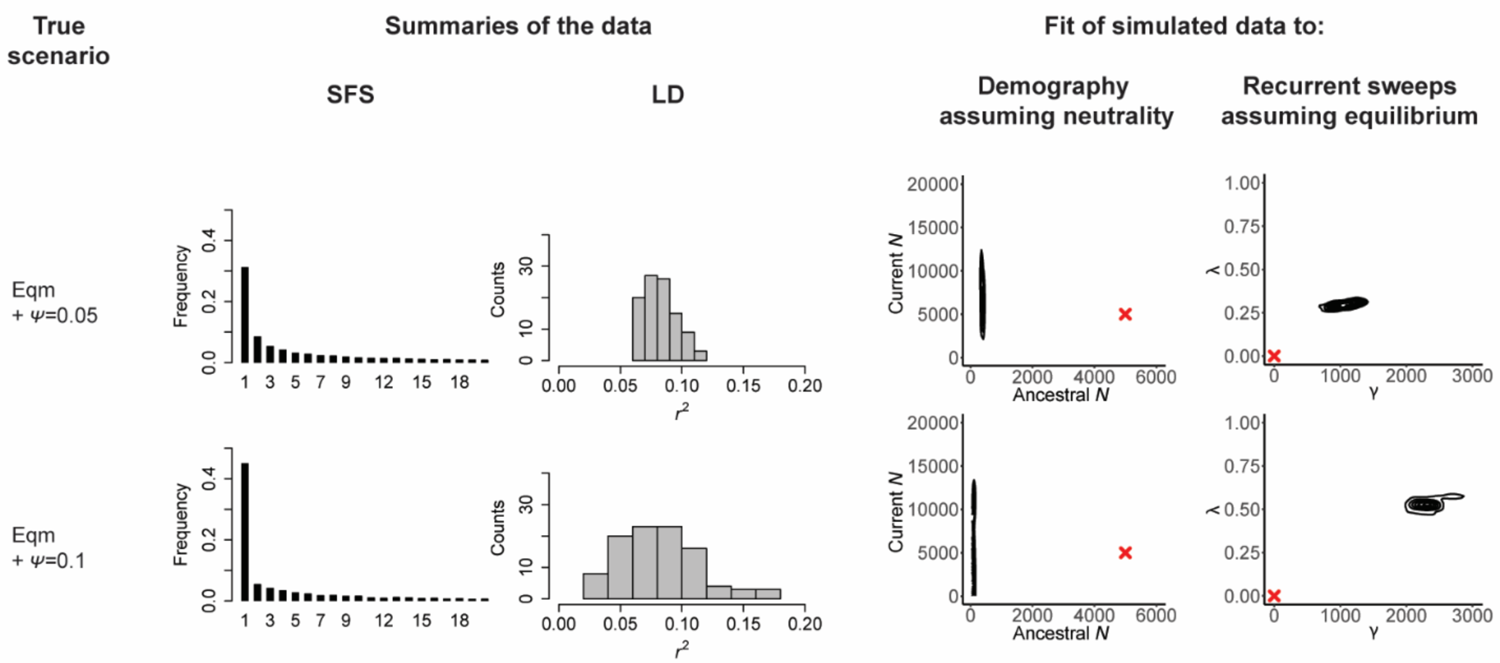
The impact of potential model violations can be quantified. As in Figures 1 and 2, the scenarios are given in the first column, here equilibrium population size together with a moderate degree of progeny skew (‘Eqm + ψ = 0.05’) as well as with a high degree of progeny skew (‘Eqm + ψ = 0.1’) (see Methodology); the middle columns present the resulting SFS and LD distributions, and the final columns provide the joint posterior distributions when the data are fit to two incorrect models: a demographic model assuming neutrality, and a recurrent selective sweep model assuming equilibrium population size. Red crosses indicate the true values. As shown, this violation of Kingman coalescent assumptions can lead to drastic mis-inference, but the biases resulting from such potential model violations can readily be described.

**Figure 4.**
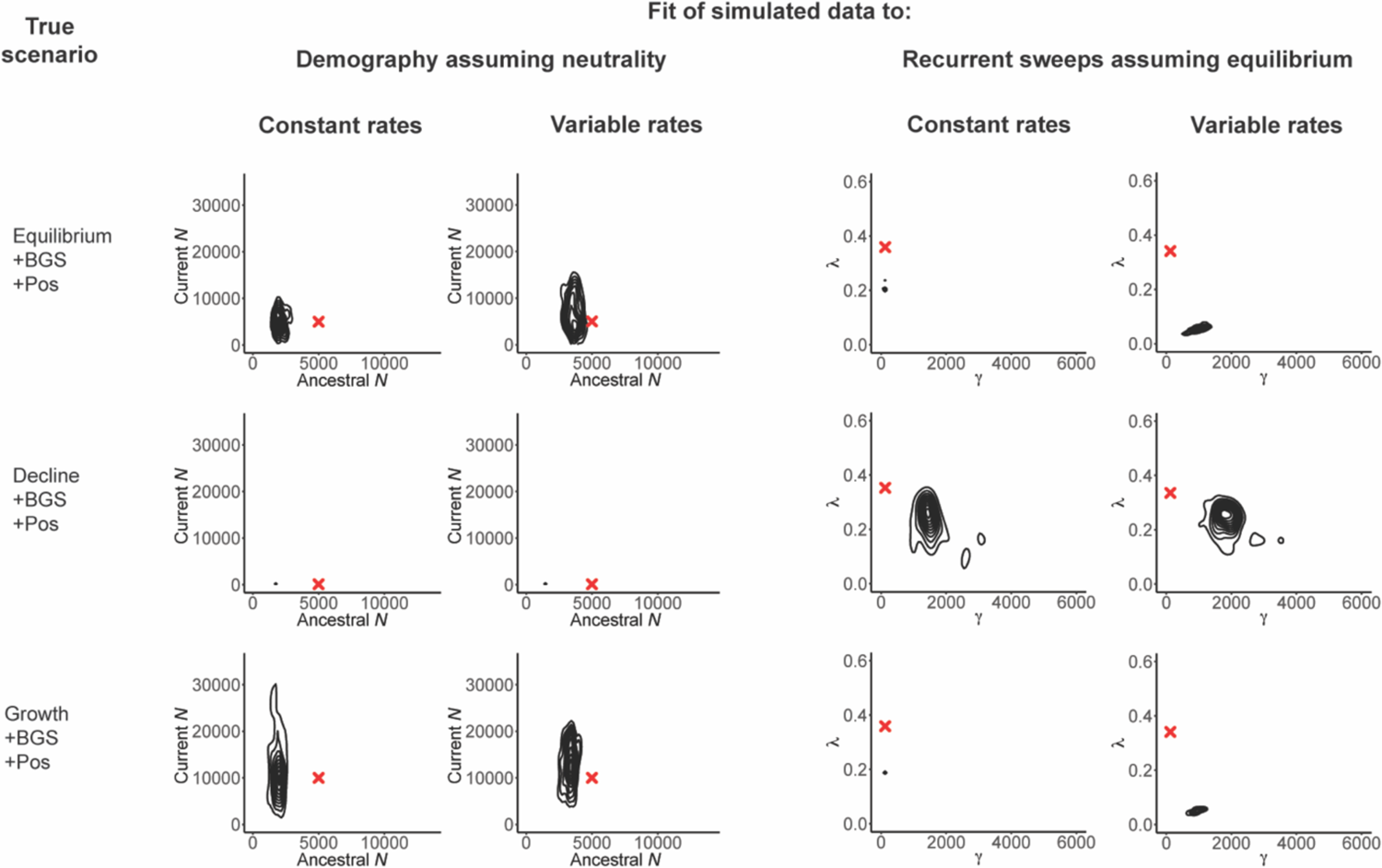
The effects of not correcting for mutation and recombination rate heterogeneity. Three scenarios are here considered: equilibrium population size with background selection and recurrent selective sweeps (‘Eqm +BGS + Pos’), declining population size together with background selection and recurrent selective sweeps (‘Decline + BGS + Pos’), and growing population size together with background selection and recurrent selective sweeps (‘Growth + BGS + Pos’). Inference is again made under an incorrect demographic model assuming neutrality, as well as an incorrect recurrent selective sweep model assuming equilibrium population size. However, within each category, inference is performed under two settings: mutation and recombination rates are constant and known, and mutation and recombination rates are variable across the region but assumed to be constant (see Methodology). Red crosses indicate the true values, and all exonic (*i.e*., directly selected) sites were masked prior to analysis. As shown, neglecting mutation and recombination rate heterogeneity across the genomic region in question can have an important impact on inference, particularly with regards to selection models.

As detailed in Figure 5, with such a model incorporating both biological and stochastic variance as well as statistical uncertainty in parameter estimates, and with an understanding of the role of likely model violations, one may investigate which additional questions / hypotheses can be addressed with the data at hand. By using a simulation approach starting with the baseline model and adding hypothesized processes, it is possible to quantify the extent to which models, and the parameters underlying those models, may be differentiated and which result in overlapping or indistinguishable patterns in the data (*e.g*., [110]). For example, if the goal of a given study is to identify recent beneficial fixations in a genome – be they potentially associated with high-altitude adaptation in humans, crypsis in mice, or drug-resistance in a virus – one may begin with the baseline model and simulate selective sweeps under that model. As illustrated in Figure 6, by varying the strengths, rates, ages, dominance and epistasis coefficients of beneficial mutations, the patterns in the SFS, LD, and/or divergence that may differentiate the addition of such selective sweep parameters from the baseline expectations can be quantified. Moreover, any intended empirical analyses can be evaluated using simulated data (*i.e*., the baseline, compared to the baseline + the hypothesis) to define the power and false-positive rates associated. If the differences in resulting patterns cannot be distinguished from the expected variance under the baseline model (in other words, if the power and false-positive rate of the analyses are not favorable), the hypothesis is not addressable with the data at hand (*e.g*., [54]). If the results are favorable, this analysis can further quantify the extent to which the hypothesis may be tested; perhaps only selective sweeps from rare mutations with selective effects greater than 1% and that have fixed within the last 0.1 *N_e_* generations are detectable (see [111–112]), and any others could not be statistically distinguished from expected patterns under the baseline model. Hence, such an exercise provides a critically essential key for interpreting the resulting data analysis.

**Figure 5:**
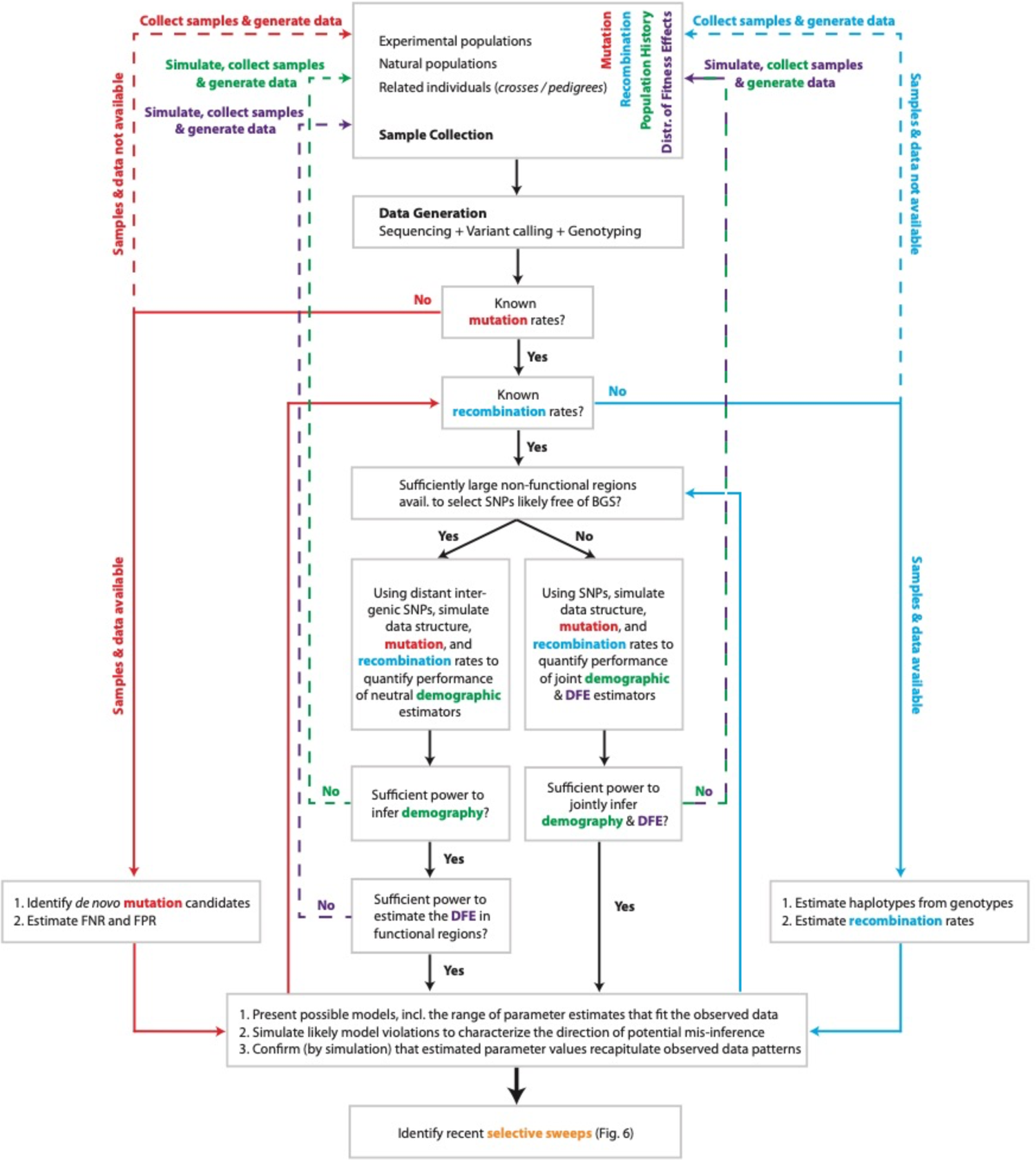
Diagram of important considerations in constructing a baseline model for genomic analysis. Considerations related to mutation rate are coded in red, recombination rate in blue, demographic history in green, and the distribution of fitness effects in purple - as well as combinations thereof. Beginning from the top with the source of data collected, the arrows suggest a path that is needed to be considered. Dotted lines indicate a return to the starting point.

**Figure 6:**
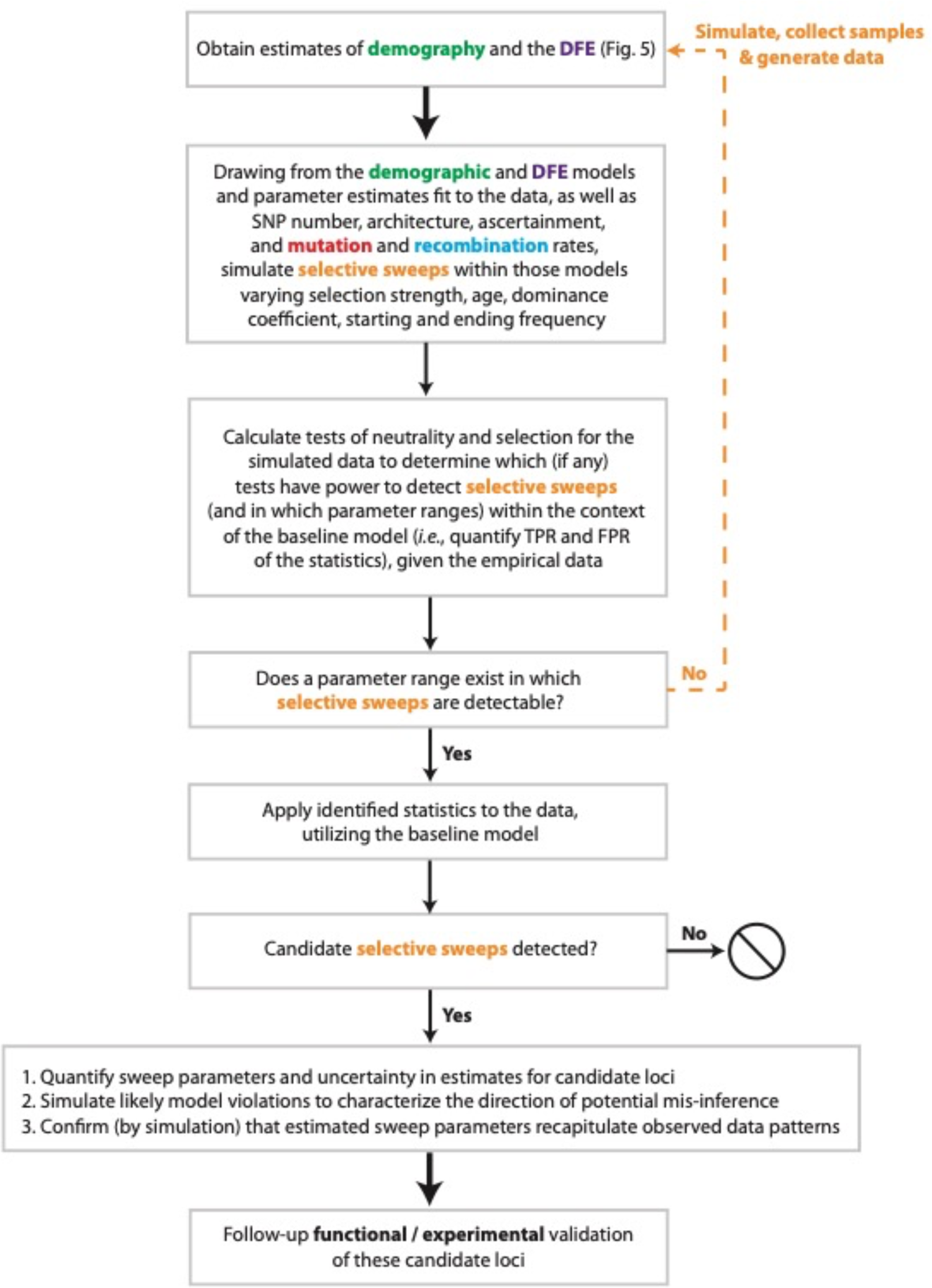
Diagram of important considerations in detecting selective sweeps. The color scheme matches that in Figure 5, with ‘selective sweeps’ coded in orange.

#### A consideration of alternative strategies

In this regard, it is worth mentioning two common approaches that may be viewed as alternatives to the strategy that we recommend. The first tactic concerns identifying patterns of variation that are uniquely and exclusively associated with one particular process, the presence of which could support that model regardless of the various underlying processes and details composing the baseline. For example, Fay & Wu’s [113] *H*-statistic, capturing an expected pattern of high-frequency derived alleles generated by a selective sweep with recombination, was initially proposed as a powerful statistic for differentiating selective sweep effects from alternative models. Results from the initial application of the *H*-statistic were interpreted as evidence of widespread positive selection in the genome of *Drosophila melanogaster*. However, Przeworski [112] subsequently demonstrated that the statistic was characterized by low power to detect positive selection, and that significant values could readily be generated under multiple neutral demographic models. The composite likelihood framework of Kim & Stephan [111] provided a significant improvement by incorporating multiple predictions of a selective sweep model, and was subsequently built upon by Nielsen *et al.* [114] in proposing the SweepFinder approach. However, Jensen *et al*. [115] described low power and high false-positive rates under certain neutral demographic models. The particular pattern of LD generated by a beneficial fixation with recombination described by Kim & Nielsen [116] and Stephan *et al*. [117] (and see [118]), was also found to be produced under an (albeit more limited) range of severe neutral population bottlenecks [119–120].

The point here is that the statistics themselves represent important tools for studying patterns of variation and are useful for visualizing multiple aspects of the data, but in any given empirical application they are impossible to interpret without the definition of an appropriate baseline model and related power and false-positive rates. Thus, the search for a pattern unique to a single evolutionary process is not a work-around, and historically such patterns rarely turn out to be process specific after further investigation. Even if a ‘bullet-proof’ test were to be someday constructed, it would not be possible to establish its utility without appropriate modeling, an examination of model violations, and extensive power / sensitivity-specificity analyses. But in reality, the simple fact is that some test statistics and estimation procedures perform well under certain scenarios, but not under others.

The second common strategy involves summarizing empirical distributions of a given statistic, and assuming that outliers of that distribution represent the action of a process of interest, such as positive selection (*e.g.,* [121]). However, such an approach is problematic. To begin with, any distribution has outliers, and there will always exist a 5% or 1% tail for a chosen statistic under a given model. Consequently, a fit baseline model remains necessary to determine whether the observed empirical outliers are of an unexpected severity, and if the baseline model together with the hypothesized process has, for example, a significantly improved likelihood.

Moreover, only by considering the hypothesized process within the context of the baseline model can one determine whether affected loci (*e.g*., those subject to recent sweeps) would even be expected to reside in the tails of the chosen statistical distribution, which is far from a given [72, 122]. As such, approaches which may not necessarily require a defined baseline model in order to perform the initial analyses (*e.g.,* [114]), nonetheless require such modeling to accurately define expectations, power and false-positive rates, and thus to interpret the significance of observed empirical outliers. For these reasons, the approach for which we advocate remains essential. As the appropriate baseline evolutionary model may differ strongly by organism and population, this performance must be carefully defined and quantified for each empirical analysis in order to accurately interpret results.

## Conclusions

When it comes to evolutionary analyses, wanting to answer a question is not necessarily equivalent to being able to answer it. The ability of population genomics to address a hypothesis of interest with a given dataset is something that must be demonstrated, and this may be achieved by constructing a model composed of common biological and evolutionary processes, including the uncertainty in those underlying parameters, as well as the specific features of the dataset at hand. The variation in possible observational outcomes associated with a chosen baseline model, and the ability to distinguish an hypothesized additional evolutionary process from such ‘background noise’, are both quantifiable. Furthermore, even if the model were correct, there exists a limit on the precision of estimation imposed by the evolutionary variance in population statistics that requires description, and which no amount of sampling can remove.

Demonstrating that multiple models, and/or considerable parameter space within a model, are compatible with the data need not be viewed as a negative or weak finding. Quite the contrary – the honest presentation of such results motivates future theoretical, experimental, and empirical developments and analyses, which can further refine the list of competing hypotheses, and this article contains many citations that have succeeded in doing this. At the same time, this analysis can define which degrees of uncertainty are most damaging (*e.g*., Figures 3 and 4), also highlighting the simple fact that organisms in which basic biological processes have been better characterized are amenable to a wider range of potential evolutionary analyses. The impact of uncertainty in these parameters in non-model organisms may motivate taking a step back to first better characterize the basic biological processes such as mutation rates and spectra via mutation-accumulation lines or pedigree studies, in order to improve resolution on the primary question of interest.

Importantly, the framework we describe will also generally identify many models and parameter realizations that are in fact inconsistent with the observed data. This ‘ruling-out’ process can often be just as useful as model-fitting, and rejecting possible hypotheses is frequently the more robust exercise of the two. The value of this narrowing down, rather than the enthusiastic promotion of individual scenarios, is worthy of heightened appreciation.

Nevertheless, all models should not be viewed equally. Decades of work supporting the central tenets of the Neutral Theory [19], high-quality experimental and computational work quantifying mutation and recombination rates [77-79, 83-84, 123], constantly improving experimental and theoretical approaches to quantify the neutral and deleterious DFE from natural population, mutation-accumulation, or directed mutagenesis data [44, 90, 124–126], and historical knowledge (*e.g*., anthropological, ecological, clinical) of population size change or structure – combined with the fact that all of these factors may strongly shape observed levels and patterns of variation and divergence – justify their role in comprising the appropriate baseline model for genomic analysis.

Given this, and particularly after accounting for the inflation of variance contributed by uncertainty in relevant parameters, potential model violations, as well as the quantity and quality of data available in any given analysis, it will often be the case that many hypotheses of interest may not be addressable with the dataset and knowledge at hand. However, recognizing that a question cannot be accurately answered, and defining the conditions under which it could become answerable, should be preferred over making unfounded and thus misleading claims.

Consistent with this call for caution however, it should equally be emphasized that the fit of a baseline model to data is certainly not inherent evidence that the model encompasses all relevant processes shaping the population. In reality, it is virtually guaranteed not to be all-encompassing, and building these models involves simplifying more complex processes (for a helpful and more general perspective, see [127]). When an additional process cannot be satisfactorily detected, that may rather be viewed as a statement about statistical identifiability – the inability to distinguish a hypothesized process from other processes that are known to be acting – and in such scenarios, absence of evidence need not be taken as evidence of absence.

While the many considerations we describe may appear daunting, it is our hope that these recommendations may serve as a useful roadmap for future data analyses in population genomics, one that may inform not only the perspectives of authors, but also that of reviewers and editors as well. Helpfully, these strategies can save considerable time, money, and effort prior to the start of empirical data handling, by determining which questions are accessible to the researcher. If a question is addressable, this preliminary analysis can additionally define what types of data are needed; for example, the number of variants or sample size necessary to obtain sufficient power, or how alternative data collections (*e.g*., temporal samples) could improve resolution. This further highlights the value of defining specific hypotheses and of studying specific patterns as opposed to running a general suite of software on each new dataset in the hopes of identifying something of interest – namely, one cannot define the power of a study to address an unformulated question. Such hypothesis-driven population genomics has resulted in a number of success stories over the past decade; systems in which specific hypotheses were formed, data was collected for the purpose, detailed population genomic analyses were designed, and ultimately important insights were gained about the evolutionary history of the population in question (*e.g*., the study of cryptic coloration has proven fruitful in this regard [3]). One feature common to these studies is interdisciplinarity: the utilization of population genetic theory and inference as described here, combined with classical genetic crosses, large-scale field studies, and genetic manipulation in order to connect genotype to phenotype to fitness and to validate statistical inference. Importantly however, without a population genetic framework for defining hypotheses, quantifying processes contributing to observed variation and divergence, evaluating and distinguishing amongst competing models, and defining uncertainty and potential biases, the observations remain merely descriptive.

## ACKNOWLEDGEMENTS

This work was funded by National Institutes of Health grants R01GM135899 and R35GM139383 to JDJ. We would like to thank Nick Barton, Matt Dean, Fabian Freund, Ryan Gutenkunst, Mark Kirkpatrick, Sarah Marion, Mohamed Noor, Sally Otto, Kevin Thornton, and John Wakeley for helpful comments and suggestions.

